# Intratumoural oncolytic HSV-1 reshapes the local and systemic immune landscape through CD8^⁺^ T cell reprogramming

**DOI:** 10.64898/2026.04.07.716986

**Authors:** Elizabeth Appleton, Anton Patrikeev, Victoria Roulstone, Mikel Portillo, Jehanne Hassan, Shane Foo, Lisa Hubbard, Joan Kyula-Currie, Isaac Dean, Ernesto Lopez, Charleen Chan Wah Hak, Malin Pedersen, Antonio Rullan, Chanidapa Tye, Mark Allen, Holly Baldock, Jessica Rowley, Emmanuel Patin, Alex Taylor, Amarin Wongariyapak, Nitya Mohan, Isla Leslie, Olivia Liseth, Benjamin Luke Kendall, Richard Vile, Esther Arwert, Sylwia Jones, Praveen Bommareddy, Robert Coffin, Masahiro Ono, Kevin Harrington, Alan Melcher

## Abstract

Most oncolytic viruses are delivered by intratumoural injection, and local administration can induce regression of both injected and distant tumours in mice and patients. However, the mechanisms by which local viral infection reprograms systemic immunity remain poorly understood. Here we show that intratumoural RP1, an oncolytic HSV-1 encoding GM-CSF and GALV-GP-R^⁻^, drives regression of injected and uninjected murine melanoma tumours and prolongs survival. RP1 elicits coordinated CD4^⁺^ and CD8^⁺^ T cell infiltration accompanied by local cytokine remodelling, reshaping the immune landscape at both tumour sites. Leveraging the Timer of Cell Kinetics and Activity (Tocky) system and Kaede photoconvertible protein technology, we resolve the temporal dynamics of CD8^⁺^ T cell responses following local virotherapy and identify two systemically induced, virus-driven CD8^⁺^ T cell populations distinguished by TCR engagement kinetics: antigen-engaged Timer-positive ‘viral-induced precursors’ (VIPs) and Timer-negative KLRG1^⁺^ ‘viral-induced terminal effectors’ (VITEs). Single-cell transcriptomic and pseudotime analyses reveal divergent differentiation trajectories; VIPs exhibit sustained antigen engagement within the tumour microenvironment (TME), a transcriptional programme associated with self-renewal, and preferential homing to draining lymph nodes. A VIP-associated gene signature correlates with clinical response to RP1 plus PD-1 blockade in the IGNYTE trial of RP1 and nivolumab in PD-1-refractory melanoma and is independently associated with response to immune checkpoint inhibitor (ICI) therapy in melanoma. These findings establish a mechanistic link between local oncolytic virotherapy, systemic CD8^⁺^ T cell reprogramming, and durable regression of distant lesions in patients.

## Introduction

Oncolytic viruses (OV) are a compelling class of immunotherapeutic agents with the unique capacity to convert immunologically “cold” tumours with sparse immune infiltration into inflamed, T cell-rich environments more amenable to effective immunotherapy^1^. By selectively replicating within and lysing tumour cells, OV simultaneously release tumour-associated antigens, stimulate innate immune sensing pathways, and, when armed with immunostimulatory transgenes, deliver localised immune payloads directly within the TME. Among the diverse OV platforms under clinical investigation, intratumoural administration of oncolytic herpes simplex virus type 1 (HSV-1) derivatives has emerged as one of the most clinically advanced strategies. Talimogene laherparepvec (T-VEC), the first OV approved by the FDA^2^, and the next-generation construct RP1 (vusolimogene oderparepvec)^3^ have demonstrated durable anti-tumour activity in a subset of patients, both as monotherapy and in combination with ICI ^4–7^. Despite this clinical progress, the immunological mechanisms that underlie these responses remain incompletely defined, particularly with respect to how local viral infection translates into systemic tumour regression.

Intratumoural administration remains the predominant delivery route for OV^8^, enabling concentrated delivery of high viral titres at the injection site while circumventing key challenges of systemic delivery including neutralising antibodies, complement activation, and pulmonary, hepatic, or splenic sequestration^2^. Local viral infection and oncolysis is conceived to trigger a cascade of immunological events, including release of tumour-associated antigens, damage-associated molecular patterns, and innate immune activation, that promote *in situ* vaccination within an immune-primed TME, leading to consequent systemic anti-tumour immunity ^9^. Although this framework has become a central paradigm in oncolytic virotherapy, direct experimental evidence supporting the model remains scarce.

Beyond the observation that intratumoural OV can induce regression of distant, uninjected lesions in a subset of patients ^10^, the mechanistic pathways by which intratumoural delivery translates into systemic immune reprogramming are largely unknown. It is not clear whether systemic reconfiguration reflects direct migration of immune cells from the injected tumour, mobilisation of a systemic reservoir, local remodelling within non-injected lesions, or a combination of these mechanisms.

RP1 is a genetically engineered HSV-1 encoding granulocyte-macrophage colony-stimulating factor (GM-CSF) and the constitutively fusogenic glycoprotein GALV-GP-R^⁻^, designed to augment both direct tumour cytolysis and immune activation^3^. RP1 represents one of the most extensively studied next-generation oncolytic HSV-1 platforms and has been shown to overcome resistance to anti-PD1 checkpoint blockade in both mice ^11^ and patients ^5^. In the phase II IGNYTE trial evaluating RP1 in combination with the anti-PD-1 antibody nivolumab in patients with checkpoint inhibitor-refractory advanced melanoma, the regimen demonstrated an overall response rate of 32.9% ^12^. Critically, in the absence of detectable RP1 spread beyond the injection site, tumour regressions occurred with comparable frequency, magnitude, and durability in both injected and distant, uninjected lesions (including visceral metastases) consistent with intratumoural RP1 priming systemic anti-tumour immunity. These findings strongly suggest that locally delivered HSV-1 can overcome resistance to checkpoint blockade and reprogram systemic anti-tumour immunity in a refractory clinical context. However, the mechanisms by which this occurs remain undefined. Understanding how the local intersection between viral and tumour immunity reprograms systemic T cell responses is essential to rationally combining OV with other immunotherapies and to identifying patients most likely to benefit.

CD8^⁺^ T cells are central mediators of tumour clearance following immunotherapy, and their differentiation from naïve to effector, progenitor exhausted (Tpex), or terminally exhausted states has been shown to critically determine the quality, magnitude, and durability of anti-tumour responses ^13,14^. Tpex CD8^⁺^ T cells have emerged as a pivotal population in cancer immunotherapy, defined phenotypically by the transcription factor TCF-1 (Tcf7) which supports self-renewal, alongside surface markers including SLAMF6 and CD62L^15^. Tpex cells maintain a self-renewing reservoir with the potential to differentiate into functional effector populations upon stimulation, and Tpex abundance and function has been consistently associated with improved responses to ICI across multiple cancer types^15–17^. This association reflects the capacity of Tpex to expand upon PD-1 blockade and fuel sustained anti-tumour effector responses, a property that terminally exhausted cells lose. In contrast, terminally exhausted CD8^⁺^ T cells are defined by loss of TCF-1, upregulation of the transcription factor TOX, and high expression of inhibitory receptors including CD39, PD-1, and TIM-3^18,19^. While exhausted T cells retain some cytotoxic function, continued exhaustion results in limited proliferative capacity and reduced or absent responsiveness to PD-1 blockade^20^ resulting in occupation of immunological space within the TME without anti-tumour activity. The accumulation of terminally exhausted CD8^⁺^ T cells is therefore frequently associated with diminished benefit from anti-PD-1 therapies and represents a major barrier to durable responses in tumours that are poorly immunogenic or that have developed resistance to ICI^20,21^.

Despite these advances, how OV therapy shapes the balance between T cell activation, Tpex maintenance, effector differentiation, and terminal exhaustion remains poorly defined. This is not a trivial gap. OV are replicating viruses that introduce dominant co-existing antiviral immunity, and the intersection between antiviral and anti-tumour T cell responses within the TME is complex and largely unexplored. The ways in which viral infection within the injected tumour influences T cell fate decisions, and how local changes propagate systemically to reshape immunity at distant sites, are open questions of direct therapeutic relevance. This is especially critical given the growing incorporation of immunomodulatory payloads into oncolytic viral backbones and the development of combination strategies that depend on exploiting local TME remodelling upon viral replication. Defining which CD8⁺ T cell populations are induced within the local environment, and how they are distributed systemically, is therefore imperative to rationally optimise therapeutic benefit.

To address this, we leveraged a syngeneic bi-flank 4434 murine melanoma mode ^22^l, a T cell-inflamed yet checkpoint-resistant BRAF^V600E^-mutant tumour representative of the clinical scenario of ICI-resistant melanoma. Using the Nr4a3-Tocky fluorescent timer reporter system^23^ alongside single-cell RNA sequencing (scRNA-seq) and paired TCR sequencing, we set out to dissect the temporal and transcriptional dynamics of local and distant CD8^⁺^ T cell responses following intratumoural RP1 administration. The Tocky transgenic system uses a fluorescent timer protein expressed downstream of the immediate-early TCR signalling gene Nr4a3 and enables quantification of real-time T cell activation dynamics *in vivo*, distinguishing recently activated from persistently antigen-engaged T cells by the ratio of blue to red fluorescence. By integrating Tocky-based temporal profiling with spatial immunophenotyping, scRNA-seq, and the Kaede^24^ photoconversion system for cellular fate-mapping, we comprehensively map how RP1 remodels the CD8^⁺^ T cell compartment at local and distant tumour sites and in secondary lymphoid organs.

This approach revealed total remodelling of the CD8^⁺^ T cell compartment in both injected and uninjected tumours upon RP1 treatment, with the emergence of two discrete virus-induced CD8^⁺^ T cell populations: terminal effector-like viral-induced terminal effectors (VITEs) which were no longer TCR engaged in the TME, and TCR-engaged progenitor exhausted-like viral-induced precursors (VIPs) which were induced bilaterally and persistently engaging with antigen within the local environment. Remarkably, these subsets exhibit distinct TCR signalling dynamics, transcriptional programmes, and differentiation trajectories, and their transcriptional signatures correlate with clinical outcomes in patients treated with RP1 plus nivolumab in the IGNYTE trial. Collectively, these findings demonstrate that intratumoural oncolytic virotherapy with RP1 orchestrates coordinated, systemic reprogramming of CD8^⁺^ T cell differentiation, establishing a mechanistic link between local viral infection and durable systemic anti-tumour immunity in both mice and patients.

## Results

### RP1 treatment induces regression of injected and distant tumours and remodels local and systemic T cell activation

Unilateral, intratumoural administration of RP1 in mice bearing bilateral 4434 melanomas led to growth inhibition of both the injected and uninjected tumours (Fig. 1a). Tumour control at both sites translated into a significant survival advantage compared with vehicle-treated animals (Fig. 1b), confirming that local delivery of RP1 is sufficient to drive systemic growth delay in this model. Notably, virus is not isolated outside the local environment in this system, by Tcid50^25^ or PCR-based assays^11^, suggesting that systemic viraemia is not driving systemic effects.

**Fig. 1.**
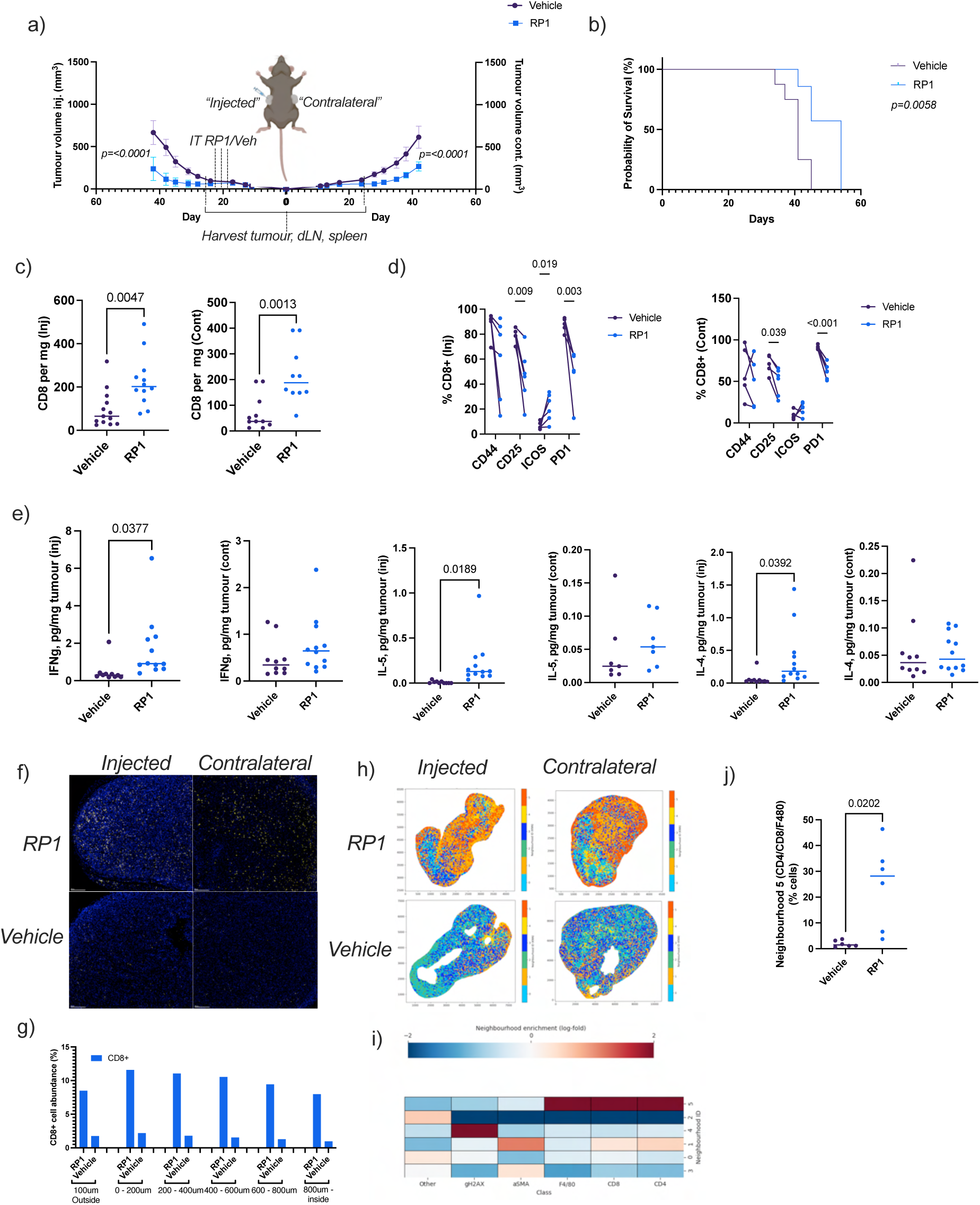
Intratumoural RP1 induces regression of injected and distant tumours and remodels local and systemic T cell architecture. **a,b,** Bi-flank 4434 melanoma-bearing mice received three intratumoural doses of RP1 (1×10^6^ PFU) or vehicle into the right-sided tumour. RP1 inhibited growth of both injected and uninjected tumours (a) and prolonged survival (b). **c,** Flow cytometric quantification of CD8^⁺^ T cells at day 7; RP1 significantly increased infiltration at both sites (p < 0.05). **d,** Activation marker expression (CD44, CD25, PD-1) was reduced in CD8^⁺^ T cells from both tumours; ICOS^⁺^ CD8^⁺^ T cells were selectively expanded in injected tumours. **e,** LEGENDplex cytokine profiling of tumour homogenates showed significant induction of IFN-γ, IL-4, and IL-5 in injected but not uninjected tumours, implicating T cell trafficking rather than systemic cytokine redistribution as the driver of distant infiltration. **f,g,** Multiplex immunofluorescence showed dense parenchyma-wide CD8^⁺^ T cell infiltration in both injected and uninjected lesions. **h–j,** Spatial neighbourhood analysis revealed RP1-induced formation of immune-enriched CD8^⁺^–CD4^⁺^–F4/80^⁺^ multicellular niches significantly expanded at both tumour sites. All flow cytometry data are representative of ≥2 independent experiments.

To evaluate how RP1 reshapes immune infiltration during the early response phase, we used an effective but non-curative treatment regimen and profiled tumours at day 7 following the first injection. Both injected and uninjected lesions displayed significantly increased infiltration of CD8^⁺^ (Fig. 1c) and CD4^⁺^ FoxP3^⁻^ T conventional cells (Supplementary Fig. 1a) relative to controls, demonstrating that viral exposure at one tumour site rapidly reconfigures the T cell landscape across injected and distant, uninjected lesions. Despite this coordinated expansion, CD8^⁺^ T cells in both tumour sites exhibited reduced expression of canonical activation and exhaustion-associated markers including CD44, CD25, and PD-1 following RP1 treatment (Fig. 1d). ICOS expression represented a notable exception, and ICOS^⁺^ CD8^⁺^ T cells were selectively increased within injected tumours, suggesting specialised costimulatory remodelling induced by RP1 at the site of viral infection ^26,27^. In contrast, CD8^⁺^ T cells from the spleens, but not lymph nodes (Supplementary Fig. 1b), of RP1-treated mice showed increased expression of CD44, CD25, PD-1, and ICOS, indicative of broad systemic activation (Supplementary Fig. 1c). These findings suggest that intratumoural RP1 drives a coordinated, systemic pattern of T cell activation, either through dissemination of secreted factors generated upon infection at the injected site, or through egress and trafficking of activated cells from the local TME.

To examine the cytokine landscape underpinning these T cell responses, we performed multi-cytokine profiling of supernatants from tumour homogenates of injected and uninjected tumours. This revealed substantial cytokine remodelling within the injected, but not uninjected tumours (Fig. 1e), including significant increases in IFN-γ, IL-4, and IL-5. The absence of cytokine changes in the uninjected tumour, despite robust T cell infiltration at that site, suggests that the coordinated T cell response in distant lesions is driven by T cell activation, differentiation, and trafficking initiated within the injected tumour and spleen, rather than by systemic cytokine dissemination from the injection site.

In addition to quantitative changes in infiltration, spatial multiplex immunofluorescence demonstrated coordinated architectural remodelling across both injected and distant tumours. RP1-treated injected lesions showed dense, parenchyma-wide CD8^⁺^ T cell infiltration relative to vehicle controls (Fig. 1f,g), and this pattern was mirrored in uninjected tumours (Fig. 1f,g), indicating that local RP1 delivery elicits disseminated, tumour-wide spatial restructuring of the immune landscape. Spatial neighbourhood analysis revealed the emergence of immune-enriched multicellular niches defined by tight spatial coupling of CD8^⁺^ T cells, CD4^⁺^ T cells, and F4/80^⁺^ macrophages (Fig. 1h,i), consistent with local opportunities for antigen presentation, T cell priming, and cytokine exchange. These organised multicellular communities were significantly expanded in both injected and uninjected lesions (Fig. 1j), demonstrating that RP1 induces not only increased immune infiltrate but also system-level rewiring of immune cell positioning and intercellular interactions at local and distant tumour sites.

### RP1 treatment triggers distinct TCR engagement states as revealed by Tocky analysis

To define how RP1 treatment reshapes early T cell activation dynamics to guide eventual cellular fate, we examined TCR signalling using the Nr4a3-Tocky reporter system in Nr4a3-Tocky mice bearing bi-flank 4434 melanoma treated with intratumoural RP1 (Fig. 2a). In the Tocky transgenic system, TCR engagement drives expression of an unstable blue, fluorescent timer protein that spontaneously and irreversibly matures to a stable red fluorescent form with a ∼4-hour half-life (Supplementary Fig. 2a)^28^. The ratio of blue to red fluorescence thus provides a unique temporal readout of TCR engagement, distinguishing newly activated (blue^⁺^), persistently engaged (blue^⁺^red^⁺^), arrested (red^⁺^) T cells *in vivo*, with timer-negative cells representing those without downstream TCR activation at the time of sampling. Representative flow cytometry gating is shown in Supplementary Fig. 2b,c.

**Fig. 2.**
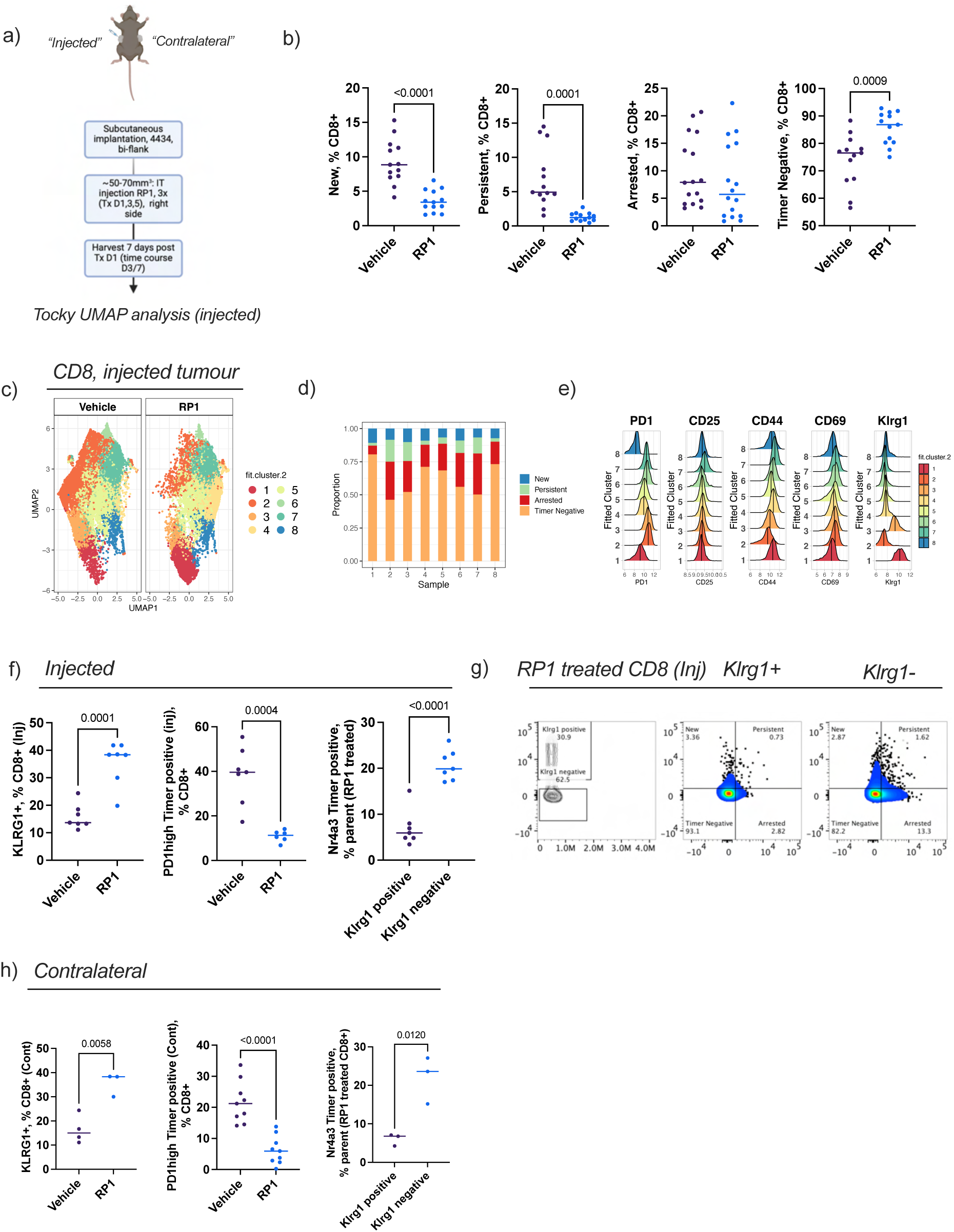
RP1 reshapes intratumoural TCR signalling dynamics in CD8⁺ T cells locally and systemically. **a,** Experimental schematic. Nr4a3-Tocky bi-flank melanoma-bearing mice treated with intratumoural RP1; tumours harvested 48 h after the final injection. **b,** RP1 caused global reduction in Tocky fluorescence in injected tumour CD8^⁺^ T cells. **c–e,** UMAP analysis revealed RP1-driven loss of PD-1^ʰʳ^ timer-positive cells (cluster 2) and induction of a KLRG1^⁺^ timer-negative population (cluster 1) alongside a KLRG1^⁻^ fraction with greater antigen engagement by Tocky. **f,g,** Representative flow plots. **h,** Analogous CD8^⁺^ T cell remodelling in the contralateral uninjected tumour, indicating systemic coordination. All flow cytometry data are representative of ≥2 independent experiments.

Across injected tumours, RP1 treatment was associated with a global reduction in Tocky fluorescence and a concomitant expansion of timer-negative CD8^⁺^ T cells (Fig. 2b). This shift reflects altered TCR engagement kinetics, potentially driven by preferential loss of pre-existing reactive clones, or dilution of timer protein driven by rapid cell division. High-dimensional UMAP analysis of CD8^⁺^ T cells revealed that this apparent global decrease in Tocky fluorescence masks profound internal restructuring of the CD8^⁺^ T cell compartment (Fig. 2c; cluster proportions in Fig. 2d; marker expression histograms in Fig. 2e). Specifically, RP1 treatment drove redistribution away from PD-1^hi^ CD44^lo^ highly timer-positive cells (cluster 2), which predominated in vehicle-treated tumours. Instead, RP1 induced two distinct populations: a timer-negative KLRG1^+^ PD-1^lo^ cluster (cluster 1) (Fig. 2c–f) and a KLRG1^⁻^ CD8^⁺^ T cell fraction that was relatively more antigen-engaged by Tocky within the RP1-treated TME (Fig. 2c–g). The KLRG1^⁺^ timer-negative phenotype is consistent with T cells that have either ceased downstream TCR signalling following effector differentiation or undergone rapid division-induced timer dilution, distinct from the KLRG1^⁻^ PD-1^int^ clusters that remained enriched for Tocky-defined TCR-engaged CD8^⁺^ T cells within the TME (Fig. 2d,f,g). As RP1 treatment reduces overall TCR-induced Tocky fluorescence, it therefore reshapes the composition of antigen-engaged CD8^⁺^ T cells, producing phenotypically distinct subsets relative to vehicle control. Strikingly, this remodelling was mirrored in the contralateral uninjected tumour (Fig. 2h), indicating that locally delivered RP1 dynamically modifies the composition of antigen-reactive CD8^⁺^ T cells at distant tumour sites.

### Single-cell transcriptomics reveals RP1-induced shifts toward precursor and cycling effector CD8⁺ T cell states with distinct TCR engagement dynamics

To further define how RP1 reshapes CD8^⁺^ T cell differentiation in relation to real-time TCR signalling, we performed scRNA-seq with paired TCR sequencing on CD8^⁺^ T cells from RP1-injected tumours, FACS sorted into the four Tocky-defined quadrants (new, persistent, arrested, and timer-negative) at day 7 following treatment (Fig. 3a,b). Following FACS, Tocky populations were barcode-labelled to enable identification in downstream single-cell analyses and association between temporal TCR signalling dynamics, transcriptional profile, and TCR clonotype.

**Fig. 3.**
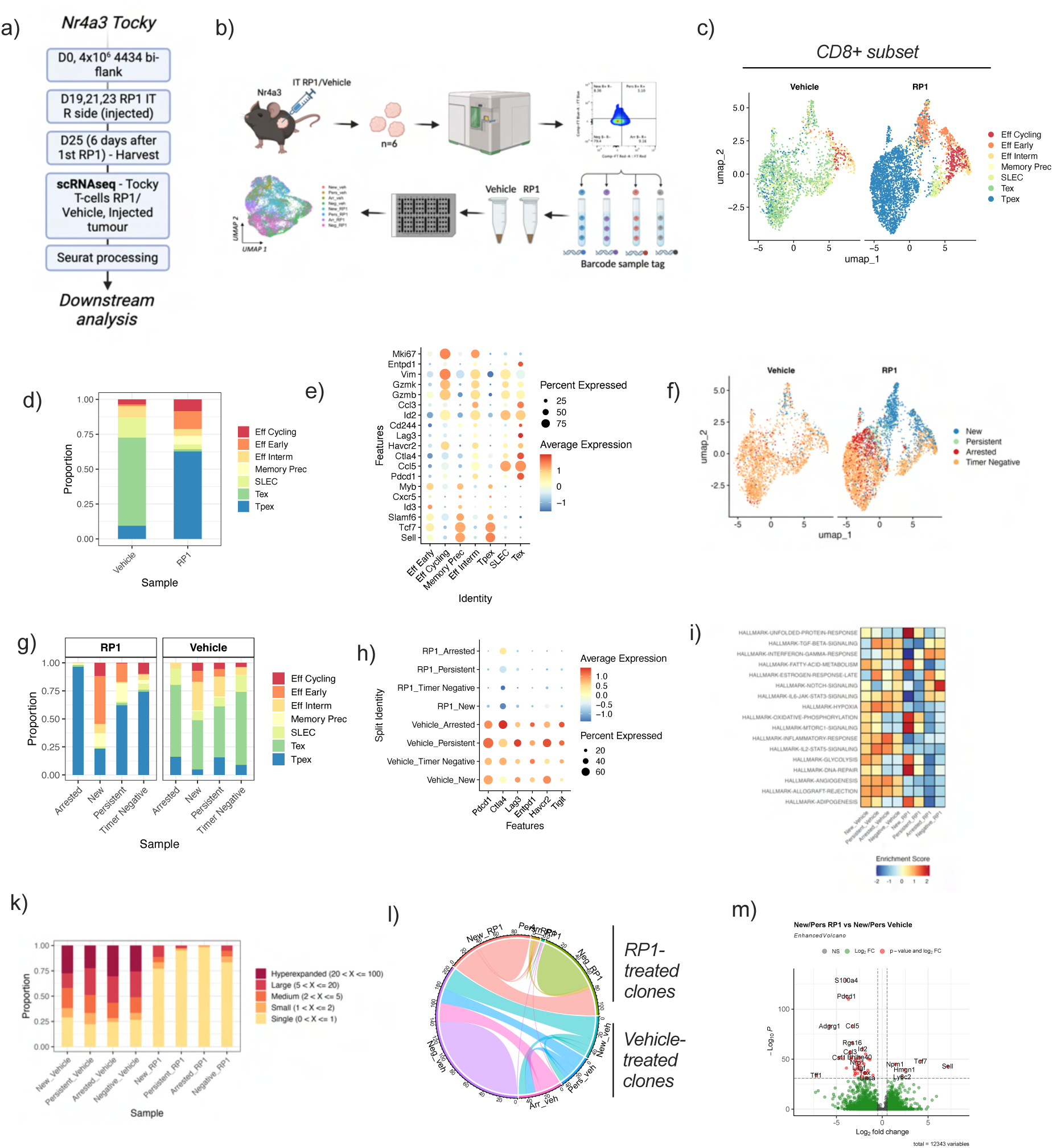
Single-cell transcriptomics reveals RP1-induced precursor and cycling effector CD8⁺ T cell states linked to distinct TCR engagement dynamics. **a,b,** Experimental schematic; CD8^⁺^ T cells from injected tumours sorted into four Tocky quadrants, barcode-labelled, and processed for scRNA-seq with paired TCR sequencing at day 6. **c,d,** Unsupervised clustering and ProjecTILs annotation showed enrichment of Tpex and cycling populations with RP1, contrasting with Tex and short-lived effector dominance in vehicle. **e,** Dot plot of marker expression by cluster. **f,g,** Tocky state mapping: persistent TCR engagement in vehicle tumours associates with Tex; in RP1-treated tumours with Tpex and memory precursors. **h,** Exhaustion marker expression by Tocky state. **i,** Hallmark pathway enrichment heatmap; RP1 upregulated DNA repair, unfolded protein response, and fatty acid metabolism. **k,l,** Reduced hyperexpanded clones and minimal vehicle/RP1 clonal overlap with RP1. **m,** Upregulation of *Tcf7*, *Sell*, *XCL1*, and *NPM1* in engaged CD8^⁺^ T cells from RP1-treated tumours.

High-resolution transcriptional profiling, UMAP hierarchical clustering, and reference-based annotation using ProjecTILs and manual lineage marker review revealed marked restructuring of the intratumoural CD8^⁺^ T cell compartment in RP1-treated mice compared with vehicle controls (Fig. 3c,d). Most notably, RP1 treatment drove enrichment of precursor-exhausted T cells expressing CD62L (*Sell*), TCF-1 (*Tcf7*), SLAMF6, and IL7R, alongside cytotoxic (expressing GZMB, GZMK, and VIM) and cycling (Ki67^⁺^) populations (Fig. 3c–e). In contrast, vehicle-treated tumours were dominated by terminally exhausted (Tex) cells expressing CTLA4, CCL5, TIM3, CD39, and PD-1, and short-lived effector cells expressing CCL5, ID2, and KLRG1 (Fig. 3c–e). These data indicate that RP1 fundamentally alters the balance of T cell differentiation states within the tumour, inducing a shift away from a dysfunctional exhaustion-dominated programme toward progenitor-enriched and proliferative states.

Mapping Tocky states onto these transcriptional clusters (Fig. 3f) revealed clear differences in antigen engagement between treatment conditions. In vehicle-treated tumours, sustained TCR engagement (Tocky persistent) was primarily associated with terminally exhausted T cells (Fig. 3g), a finding consistent with prior reports linking chronic TCR stimulation to terminal exhaustion in the tumour setting. In RP1-treated tumours, by contrast, sustained TCR engagement was primarily associated with Tpex and memory precursor cell states (Fig. 3g). Notably, TCR engagement within the virus-treated TME was associated with a global reduction in exhaustion marker expression (Fig. 3h), demonstrating that cells within the RP1-conditioned environment are able to sustain TCR signalling without exhaustion. This links sustained TCR signalling within a viral cytokine-conditioned TME to Tpex maintenance rather than terminal exhaustion, a potentially critical distinction for subsequent responsiveness to PD-1 blockade. Pathway analysis further revealed RP1-induced upregulation of DNA repair pathways, unfolded protein response signatures, and metabolic programmes including fatty acid metabolism (Fig. 3i), features associated with enhanced T cell proliferative fitness and long-term persistence.

Paired TCR analysis revealed further divergence between treatment groups. RP1-treated tumours showed increased TCR diversity across all Tocky quadrants, with fewer large or hyperexpanded clones relative to vehicle (Fig. 3k). RP1 treatment induced near-complete clonal replacement, with minimal overlap in dominant clonotypes between RP1-treated and vehicle conditions (Fig. 3l). These data indicate that RP1-mediated recruitment of CD8^⁺^ T cells reshapes the intratumoural TCR repertoire, replacing dominant dysfunctional clones with a diverse population of precursor-like, antigen-engaged T cells. The breadth of this repertoire suggests that, although expanded in the context of viral therapy, these T cells are unlikely to be exclusively HSV-specific and represent a source for propagation of anti-tumour immunity upon subsequent checkpoint blockade.

Differential expression analysis focusing on recently TCR-engaged cells (new and persistent Tocky quadrants) demonstrated marked upregulation of precursor- and memory-associated genes including *Tcf7* (TCF-1), *Sell* (CD62L), *Xcl1*, and *Npm1* in recently engaged CD8^⁺^ T cells from RP1-treated tumours compared with vehicle controls (Fig. 3m). These data indicate that CD8^⁺^ T cells experiencing active or sustained TCR signalling within the RP1-treated TME preferentially adopt progenitor-like, self-renewing states rather than the terminal exhaustion that characterises antigen-engaged cells in vehicle-treated tumours.

Using the ProjecTILs reference atlas ^29^ integrating both viral (LCMV) and tumour-infiltrating lymphocyte (TIL) datasets, we annotated our single-cell data to resolve transcriptomic differences between RP1-induced precursor populations and classical tumour-infiltrating Tpex cells. CD8^⁺^ T cells were mapped onto TIL and LCMV reference landscapes (Supplementary Fig. 4a), revealing that cells from RP1-treated tumours predominantly align with viral Tpex states, whereas cells from vehicle-treated tumours map closely to exhausted (Tex) populations regardless of the atlas used for annotation. These findings underscore fundamental differences between tumour- and virus-associated immune states within the intersecting immunological context of oncolytic virotherapy in cancer. Differential gene expression further showed that viral Tpex cells exhibit increased expression of self-renewal markers (*Tcf7*, *Slamf6*) alongside elevated cytokine and chemokine genes (*Il4i1*, *Cxcl10*) compared with tumour Tpex from the reference atlas (Supplementary Fig. 4c), pointing to a distinct, potentially more plastic precursor state induced by the viral inflammatory context.

### Single-cell transcriptional and trajectory analysis identifies two RP1-induced CD8⁺ T cell lineages with distinct TCR engagement dynamics

To resolve the differentiation programmes induced by RP1 more precisely, we focused single-cell analyses on TCR-engaged, non-naïve CD8^⁺^ T cells isolated from RP1-injected tumours, FACS sorted and labelled by Tocky quadrant. Unsupervised clustering and UMAP projection (Fig. 4a) revealed two major transcriptional clusters, each composed of multiple linked states but separating in part by their TCR signalling dynamics and beginning with new TCR engagement as defined by Tocky (Fig. 4b).

**Fig. 4.**
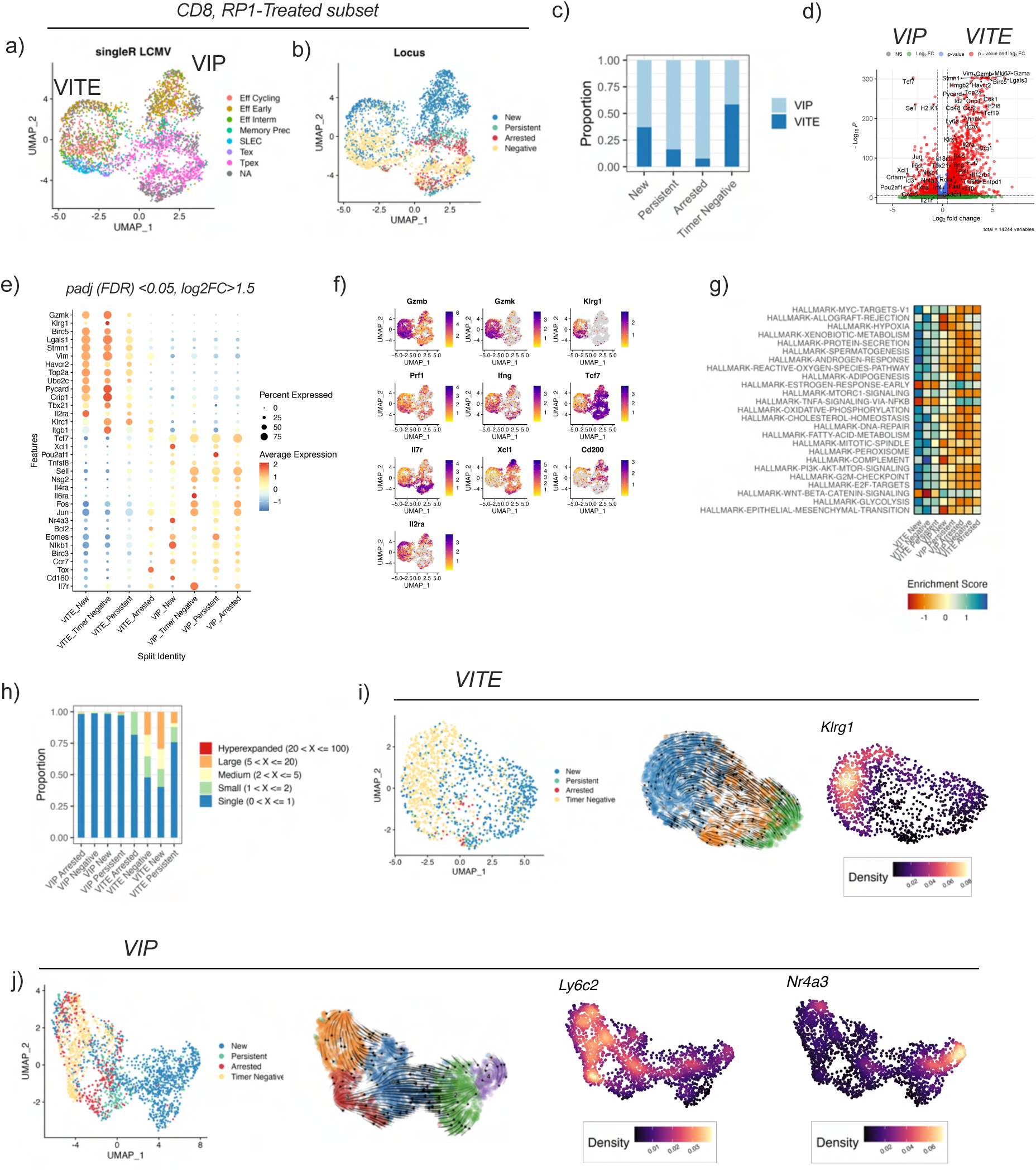
RP1 induces two transcriptionally and functionally distinct CD8⁺ T cell lineages with divergent TCR engagement dynamics. **a,** UMAP of non-naïve, TCR-engaged CD8^⁺^ T cells from RP1-injected tumours sorted by Tocky quadrant; two major transcriptional clusters identified. **b,c,** Tocky overlay: one cluster enriched for timer-new/negative cells (VITEs); second spanning new, persistent, and arrested states (VIPs). **d–f,** VITEs express KI67, GZMB, KLRG1, ENTPD1, BIRC5, and VIM; VIPs express *Tcf7*, *Sell*, *JUN*, *XCL1*, *Nr4a3*, *Il4ra*, and *Il6ra*. **g,** VITEs enriched for cytotoxicity and glycolysis; VIPs for immune homeostasis, metabolic flexibility, and transcriptional plasticity. **h,** Clonal expansion in VITEs versus diverse low-frequency clones in VIPs. **i,j,** VITE trajectories progress from timer-new to timer-negative with distal KLRG1; VIP trajectories span timer-new through persistent to arrested, with early *Nr4a3* and late *Ly6c*.

The first cluster consisted largely of timer-new and timer-negative cells. Differential expression analysis demonstrated transcriptional alignment with a KLRG1^⁺^ terminal effector-like state, expressing Ki67, Gzmb, Klrg1, Entpd1 (CD39), Birc5, and Vim (Fig. 4d–f). These cells, which we refer to as viral-induced terminal effectors (VITEs), exhibited minimal ongoing antigen engagement within the TME by Tocky, consistent with rapid effector differentiation, timer protein dilution through proliferation, or cessation of downstream TCR signalling following terminal effector commitment. The majority of persistent TCR signalling within the TME was instead found within the second cluster (Fig. 4b,c), consistent with our flow cytometry data showing that timer-positive cells persist within the RP1-treated TME despite a global reduction in Tocky fluorescence. This second cluster displayed transcriptional features of progenitor-like differentiation, including *Sell* (CD62L), *Tcf7* (TCF-1), *Jun*, and *Xcl1* (Fig. 4d–f), alongside TCR signalling-related transcripts including *Nr4a3* itself (Fig. 4e). These viral-induced precursor cells (VIPs) demonstrated sustained or arrested TCR signalling within the TME as identified by Tocky (Fig. 4c), and additionally upregulated cytokine-response receptors including *Il4ra* and *Il6ra* (Fig. 4d,e,g), consistent with a microenvironmental context that promotes precursor maintenance and responsiveness to homeostatic cytokine signals.

Hallmark gene signature and pathway analysis revealed deeply divergent biological programmes between the two lineages (Fig. 4g). VITEs were enriched for cytotoxic effector and glycolytic pathways, consistent with a short-lived effector programme. VIPs, by contrast, showed signatures associated with immune homeostasis, oxidative and fatty acid metabolism, stress adaptation, and transcriptional plasticity, features consistent with progenitor potential and long-term persistence ^30,31^. TCR repertoire analysis further supported functionally distinct roles for the two populations (Fig. 4h). VITEs demonstrated evidence of clonal expansion, consistent with their effector-biased state. In contrast, VIPs remained largely composed of diverse, low-frequency clones despite their antigen-engaged Tocky profiles, suggesting this precursor pool is sustained by ongoing recruitment into the tumour rather than by expansion of dominant, pre-existing clones at this timepoint.

Trajectory inference using Slingshot and RNA velocity analysis aligned with the temporal information encoded by Tocky (Fig. 4i,j). VITE trajectories followed a progression from timer-new to timer-negative states, consistent with rapid proliferation, timer protein dilution, and short-lived effector differentiation, with enrichment of Klrg1 expression towards the distal end of the trajectory (Fig. 4i). VIP trajectories spanned timer-new to persistent to arrested states, reflecting continued antigen engagement and a progenitor-like differentiation path, with early expression of *Nr4a3* and late expression of *Ly6c* (Fig. 4j).

Collectively, these data reveal that RP1 induces two functionally and transcriptionally distinct CD8^⁺^ T cell lineages whose identity is tightly linked to their TCR signalling dynamics within the TME. Tpex cells are classically viewed as a minor fraction of tumour-infiltrating T cells, yet they are a well-established and critical component of the acute antiviral immune response. Here, we demonstrate a substantial Tpex subset that is induced by RP1 and that is preferentially activating gene expression downstream of TCR signalling within the TME therefore likely engaging with locally presented antigens. Viral-induced inflammation and cytokine conditioning may thus support infiltration of diverse precursor subsets that subsequently engage with antigens displayed in the TME, creating a pool of antigen-experienced Tpex primed for expansion upon PD-1 blockade.

### RP1 drives systemic reshaping of the CD8⁺ T cell landscape and promotes migration of precursor-like cells to secondary lymphoid organs

To determine whether RP1-induced CD8^⁺^ T cell programmes extend beyond the injected tumour, we performed scRNA-seq with paired TCR sequencing on CD8^⁺^ T cells isolated from both injected and uninjected tumours from RP1-treated bi-flank 4434 melanoma-bearing mice (Fig. 5a). Unsupervised clustering (Fig. 5b) demonstrated restructuring of CD8^⁺^ T cell populations with striking concordance across both tumour sites in RP1-treated mice, with bilateral enrichment of Tpex-like precursor cells and cycling effector states mirroring the populations identified through Tocky-enriched scRNA-seq analyses of the injected tumour (Fig. 5c). Paired TCR sequencing confirmed that transcriptional changes were accompanied by increased TCR diversity in both injected and uninjected lesions (Fig. 5d).

**Fig. 5.**
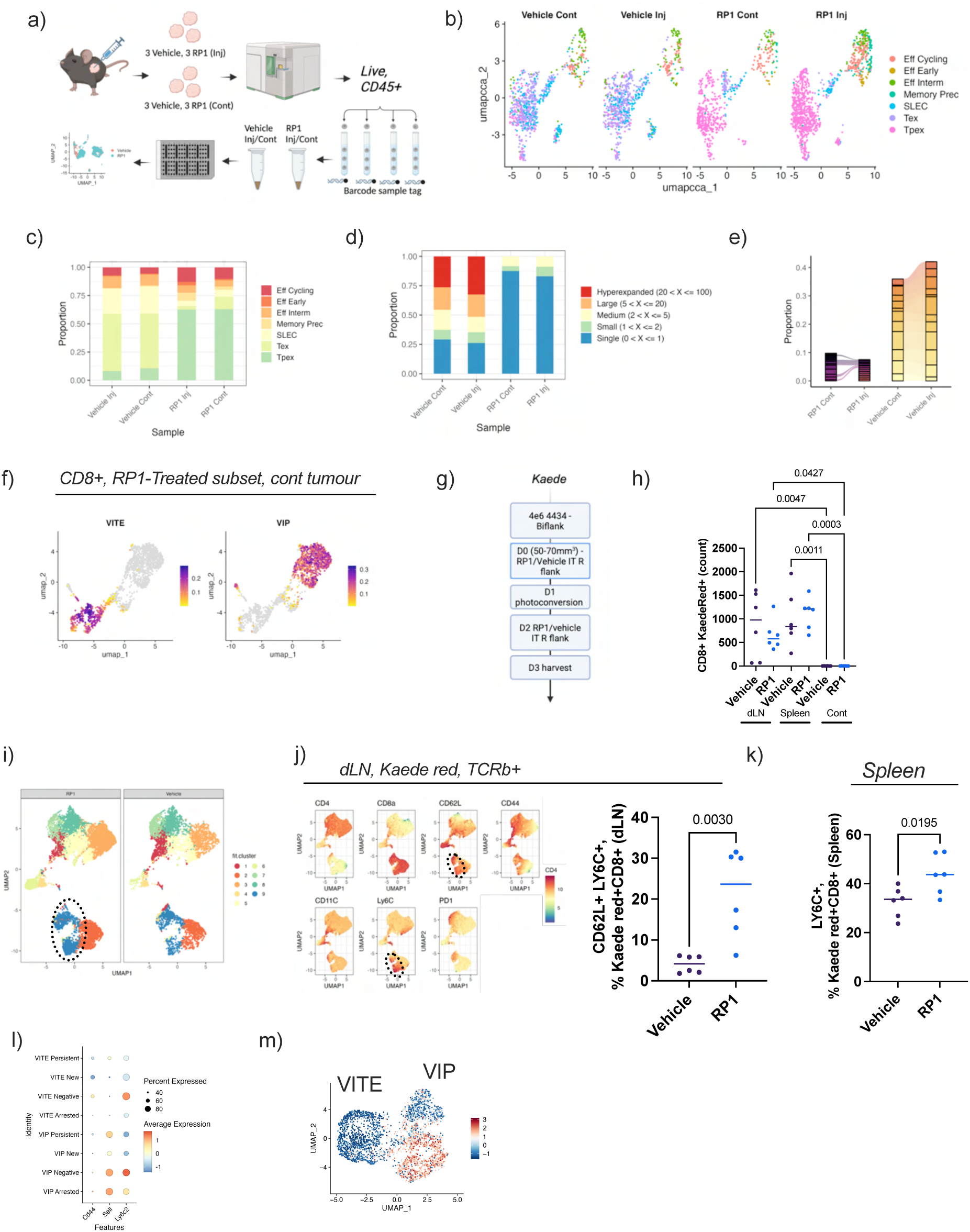
RP1 drives systemic CD8⁺ T cell reshaping and promotes precursor egress to lymphoid organs. **a,** Experimental schematic. **b,c,** scRNA-seq from injected and uninjected tumours: bilateral enrichment of Tpex-like precursors and cycling effectors with RP1. **d,** Increased TCR diversity at both tumour sites in RP1-treated mice. **e,** Vehicle-treated mice share dominant TCR clones across tumour sites; RP1-treated mice show non-overlapping, diverse repertoires at each site. **f,** VIP and VITE module scores confirm bilateral induction of both lineages. **g,h,** Kaede fate-mapping: injected tumour photolabelled after first RP1 dose; tissues harvested 48 h later. Kaede-red CD8^⁺^ T cells in tumour-draining lymph nodes and spleen but not uninjected tumours, indicating that direct tumour-to-tumour migration is not the primary mechanism of distant remodelling at this timepoint. **i,j,** RP1-induced enrichment of Ly6C^⁺^ CD62L^⁺^ cells (cluster 9) in draining lymph node Kaede-red CD8^⁺^ T cells. **k,** Corresponding increase in Ly6C^⁺^ CD8^⁺^ T cells in the spleen. **l,m,** Ly6c and Sell enrichment at distal VIP trajectory, consistent with precursor transition toward a migratory lymph node-homing phenotype.

Strikingly, TCR repertoires diverged markedly between treatment groups. In vehicle-treated mice, injected and uninjected tumours shared a substantial fraction of TCR clones (Fig. 5e), consistent with coordinated expansion of common tumour-reactive clones that were nonetheless insufficient for tumour control. In RP1-treated mice, bilateral clonal overlap was markedly reduced (Fig. 5e); instead, RP1 was associated with distinct, highly diverse TCR repertoires at injected and uninjected tumour sites. These findings suggest that RP1 drives recruitment or activation of diverse T cell populations independently within each tumour microenvironment, rather than systemic expansion of a shared dominant clone. An alternative, non-mutually exclusive interpretation is that RP1 selectively eliminates or suppresses dysfunctional, hyperexpanded T cell clones at both sites, releasing immunological space and enabling broader antigenic coverage in place of a response skewed toward a limited set of ineffective clones. The breadth of the induced repertoire is inconsistent with exclusively HSV-specific T cell priming and supports the conclusion that RP1 re-seeds both tumour sites with diverse, potentially tumour-reactive T cell populations.

Application of VIP and VITE module scores derived from the injected tumour analyses confirmed that RP1 induces discrete precursor-like and terminal effector-like states in the uninjected tumour, recapitulating the bifurcation observed in injected lesions (Fig. 5f). The two virus-associated lineages are therefore systemically induced and are not restricted to the locally injected tumour.

To investigate whether trafficking of tumour-experienced T cells contributes to systemic reshaping of the distant TME following local therapy given the absence of detectable virus and significant cytokine remodelling in the uninjected tumour (Fig. 1e), we used the Kaede UV photoconversion model^24,32^ to fate-map CD8^⁺^ T cells originating from the injected tumour. The Kaede system uses transdermal UV photolabelling to irreversibly convert the fluorescence of cells within the local TME at the time of labelling from green to red (Supplementary Fig. 3a), enabling subsequent tracking of labelled cells at distant sites by flow cytometry. Kaede mice bearing bi-flank 4434 melanomas were treated with intratumoural RP1; the injected tumour was photoconverted the day after the first RP1 dose and a further dose of RP1 was delivered into the injected tumour 48 hours later (Fig. 5g). At 48 hours post-photolabelling, approximately 50% of CD45^⁺^ cells (Supplementary Fig. 3c) and 20% of CD8^⁺^ T cells (Supplementary Fig. 3d) within the injected tumour were Kaede red.

Kaede-red CD8^⁺^ T cells were detected in tumour-draining lymph nodes and spleen, but not in uninjected tumours (Fig. 5h), indicating that direct migration from the injected tumour to a distant tumour site is not the primary source of T cell remodelling of distant lesions at this timepoint.

UMAP analysis of gated Kaede-red CD8^⁺^ T cells from the injected tumour draining lymph node showed a marked RP1-induced increase in CD8^⁺^ Ly6C^⁺^ CD62L^⁺^ cells (cluster 9, corresponding to the distal end of the VIP trajectory in our single-cell dataset) (Fig. 5i,j), with a corresponding increase in Ly6C^⁺^ CD8^⁺^ T cells in the spleen (Fig. 5k). This suggests that CD62L^⁺^ Ly6C^⁺^ CD8^⁺^ T cells are therefore preferentially egressing from the injected tumour and trafficking to secondary lymphoid organs at an early timepoint following RP1 treatment. Overlaying Ly6c and *Sell* expression as a module score onto the injected tumour single-cell dataset (Fig. 5l,m) revealed enrichment of these markers at the distal end of the VIP trajectory, consistent with precursor-like cells transitioning away from sustained antigen engagement within the TME toward a migratory lymph node-homing phenotype. Together, these data support a mechanistic model in which RP1 induces a diverse pool of antigen-engaged CD8^⁺^ precursor cells that sustain TCR signalling within the TME, then exit the tumour to establish a reservoir in secondary lymphoid tissues, providing a platform from which antigen-reactive T cells can disseminate to uninjected tumour sites and mediate systemic anti-tumour immunity.

### VIP-associated transcriptional programmes correlate with clinical response to RP1 plus PD-1 blockade and to immune checkpoint therapy in melanoma

To assess the translational relevance of RP1-induced CD8^⁺^ T cell states identified in mice, we analysed bulk RNA-seq from paired tumour biopsies from ICI-refractory melanoma patients enrolled in the IGNYTE study (NCT03767348), obtained pre-treatment and on-treatment at day 43 (prior to cycle 4) during therapy with intratumoural RP1 and PD-1 blockade (Fig. 6a). Response was assessed by RECIST criteria; notably, and consistent with the systemic immune reprogramming identified in our murine studies, responses in IGNYTE were observed with comparable frequency, magnitude, and durability in both injected and uninjected lesions ^5^.

**Fig. 6.**
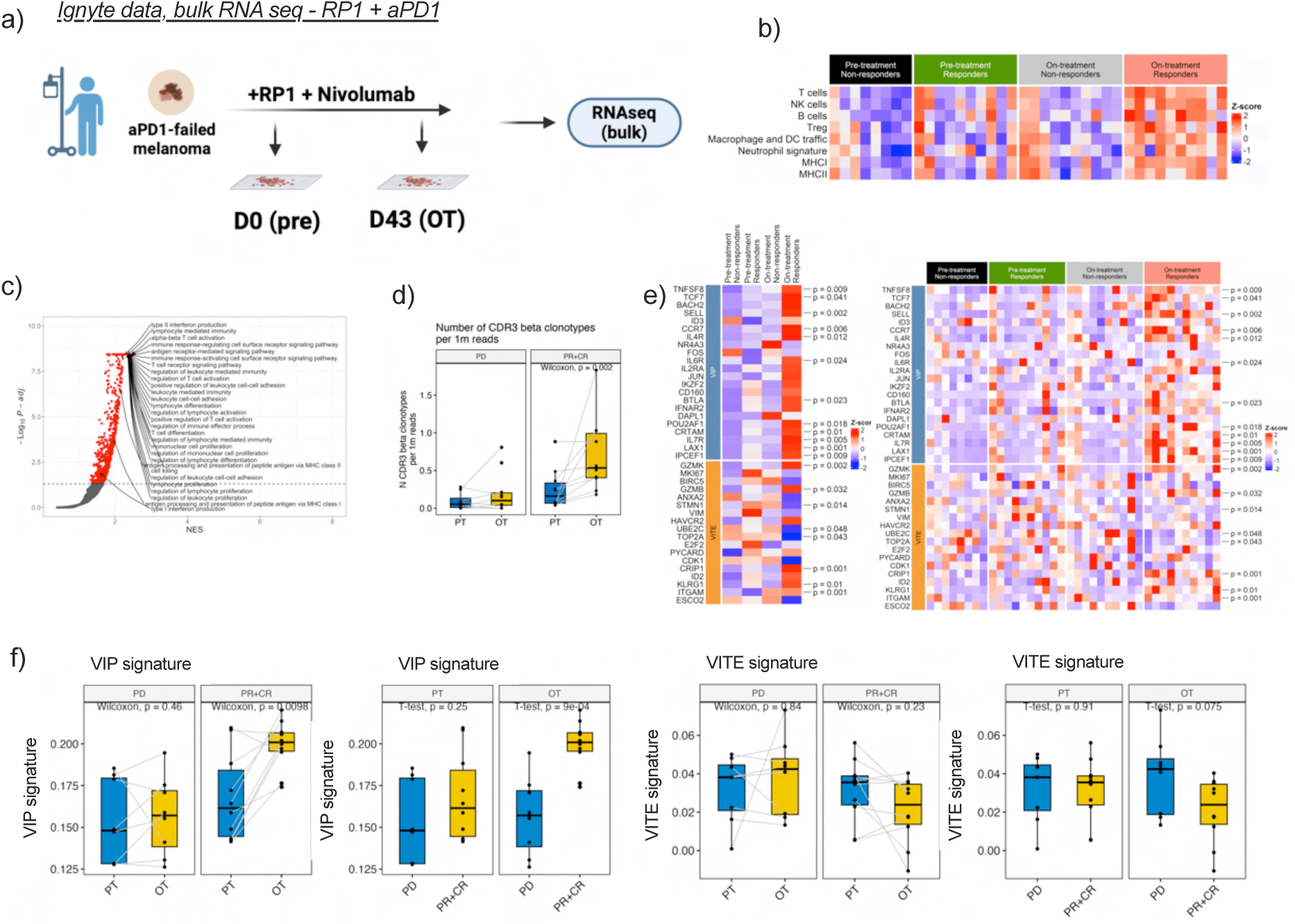
VIP transcriptional programmes correlate with clinical response to RP1 plus PD-1 blockade and to ICI in melanoma. **a,** Paired FFPE biopsies from ICI-refractory melanoma patients in IGNYTE (NCT03767348) obtained pre-treatment and on-treatment (day 43). **b,** Bulk RNA-seq deconvolution: induction of diverse immune infiltrate in responders. **c,** Enrichment of T cell activation pathways in responders. **d,** Increased TCR diversity in responders by MiXCR. **e,** VIP-associated genes (TCF7, SELL) significantly enriched in on-treatment biopsies from responders; VITE genes unchanged. **f,** Composite VIP gene signature associated with clinical response; VITE signature showed no association. **g,** Publicly available scRNA-seq from melanoma patients receiving primary ICI: VIP but not VITE enrichment in CD8^⁺^ T cells from responding tumours. Kaplan–Meier analysis of the GIDE cohort: improved overall survival in responders with high VIP scores.

Deconvolution of bulk RNA-seq data demonstrated clear induction of diverse immune infiltrate in responding patients (complete plus partial responders) (Fig. 6b). Differential gene expression analysis comparing responders and non-responders revealed enrichment of T cell activation-associated pathways in responders (Fig. 6c), with increased TCR diversity in responders by MiXCR deconvolution of bulk RNA-seq data (Fig. 6d), mirroring the increased TCR diversity observed in RP1-treated murine tumours. Analysis of viral-induced signature enrichment by response showed significant enrichment of VIP-associated genes in responding patients in on-treatment samples at both group and individual patient level (Fig. 6e), including TCF7 and SELL, mirroring the precursor-like transcriptional programmes enriched in RP1-injected mouse tumours. In contrast, these transcripts remained low or unchanged in non-responders. A composite VIP gene signature was significantly enriched through treatment and associated with clinical response in on-treatment samples from IGNYTE patients (Fig. 6f), whereas the VITE signature showed no association with therapeutic outcome. These findings indicate that expansion or preservation of diverse, precursor-like CD8^⁺^ T cell transcriptomic states, rather than terminal effector programmes, characterises tumours that respond to RP1 plus PD-1 blockade in this ICI-refractory cohort.

To determine whether this relationship extends beyond RP1 therapy, we analysed publicly available scRNA-seq datasets from melanoma patients receiving immune checkpoint inhibitors for the first time^33^. Responding tumours demonstrated enrichment of the VIP signature in CD8^⁺^ T cells sampled during treatment (Supplementary Fig. 5a), consistent with the established association between Tpex-like states and responsiveness to PD-1 blockade in melanoma. The VITE signature was not enriched in responders (Supplementary Fig. 5a). Assessment of the VIP signature within the GIDE melanoma cohort further demonstrated improved overall survival in responding patients with high VIP scores (Supplementary Fig. 5b). Together, these analyses reveal that the precursor-like transcriptional state induced by RP1 in mice has direct and significant clinical relevance: VIP-like transcriptional programmes emerge in human tumours during effective RP1 plus PD-1 treatment, and are characteristic of successful responses to primary ICI, and are associated with improved survival. These data link the effects of local viral therapy to systemic anti-tumour T cell reprogramming and to clinical benefit from OV-ICI combinations which may guide rational therapeutic design.

## Discussion

These data demonstrate that intratumoural administration of the oncolytic HSV RP1 elicits a coordinated reshaping of the CD8^⁺^ T cell compartment that extends beyond the injected lesion, presenting a mechanism for epitope spreding and priming of systemic anti-tumour immunity. It is important to recognise that OVs are not purely immunotherapies but replicating viruses that introduce dominant co-existing antiviral immunity. While prior studies have begun to characterise OV-induced responses against viral epitopes^19^, the emergence of diverse precursor populations described here represents an additional, beneficial, and until now poorly characterised component of the OV-induced immune response. Here, we show that such populations arise both within the injected tumour and systemically and are differentially associated with response to RP1-aPD1 therapy in refractory melanoma patients, establishing a new biological model for how local oncolytic virotherapy can propagate durable systemic anti-tumour immunity.

We observe systemic remodelling of T cell states characterised by the emergence and maintenance of precursor-like CD8^⁺^ T cells that retain active antigen engagement within the TME, as demonstrated using the Tocky *in vivo* system. These VIP cells, defined by sustained TCR signalling and a Tpex-like transcriptional state, appear to engage within the local inflammatory milieu generated by viral infection. Such prolonged antigen exposure within a favourable cytokine-conditioned TME may create an opportunity for priming or re-priming of this diverse influx of precursor cells against tumour antigens liberated by viral oncolysis. Tpex cells have been well established as an integral population for effective response to anti-PD-1 therapy, with demonstrated stem-like properties and capacity to expand upon checkpoint blockade into functional effector cells^9^. Prior research has shown that Tpex represent a minor fraction of tumour-infiltrating T cells outside the context of OV therapy, and optimal TCR engagement has been identified as a key determinant of Tpex maintenance^19^. Here, using Tocky, we demonstrate that Tpex are abundant within local and distant TME upon intratumoural RP1 treatment, are associated with sustained TCR signalling, and carry a transcriptomic signature that is associated with clinical response to RP1 plus anti-PD-1 in refractory melanoma patients.

Clonal analyses indicate that RP1 does not simply amplify pre-existing clones. Instead, treatment promotes increased TCR diversity across local and distant tumour sites and repopulation by distinct clones locally and remotely, displacing the exhausted, hyperexpanded repertoire that hallmarks ineffective chronic T cell responses. The lack of shared clones between injected and uninjected tumours in RP1-treated mice, in contrast to vehicle controls where dominant clones are shared between sites, supports a paradigm in which RP1 drives local antigen engagement and establishes a diverse Tpex reservoir for subsequent checkpoint blockade and expansion of subdominant tumour-reactive clones, rather than reinvigoration of an exhausted dominant repertoire. This diversification at both tumour sites, together with precursor enrichment, represents a fundamentally different immunological state from the dysfunctional, clonally restricted T cell responses that characterise untreated or vehicle-treated tumours.

In the Kaede photoconversion model, precursor-like Ly6C^⁺^ CD62L^⁺^ CD8^⁺^ T cells preferentially traffic to lymphoid tissues following prolonged antigen engagement within the tumour. This migratory behaviour provides a plausible route for antigen transport, epitope spreading, and priming of T cells outside the TME, reinforcing the capacity of RP1 to reshape systemic immunity rather than merely amplify local effector responses. RP1 therefore does not rely solely on direct lesion-to-lesion dissemination of T cells to drive distant responses; instead, it induces systemic remodelling of the CD8^⁺^ T cell compartment in secondary lymphoid organs through preferential trafficking of VIP cells from the injected tumour, providing a reservoir that may re-seed both uninjected tumours and circulating immunity.

One unresolved question concerns the antigen specificity of CD8^⁺^ T cell clones emerging during the RP1 response. Our working model is that expanded VITE clones represent an early antiviral short-lived effector response that remodels the TME alongside the emergence of diverse Tpex-like VIPs, which are given the opportunity to recognise tumour antigens, as evidenced by downstream TCR signalling within this subset detected by Tocky, and subsequently expand upon ICI. We are actively investigating TCR specificity through tetramer assays and functional testing of identified TCRs against candidate viral and tumour antigens.

The translational relevance of our murine findings is strongly supported by patient data from IGNYTE. Responders exhibit increased expression of VIP-associated genes in on-treatment tumour biopsies, increased TCR diversity, and significant enrichment of the composite VIP gene signature, while VITE signatures are not associated with response. Strikingly, the VIP signature also predicts response to ICI independently of RP1 in treatment-naïve patients both pre- and on-treatment, and associates with improved survival. This suggests that RP1 reconditions the TME in ICI-resistant patients by restoring the Tpex-aligned CD8^⁺^ T cell state that is a prerequisite for successful checkpoint blockade. These data highlight the relevance of Tpex-like cell states in OV therapy (aligned with other facets of immunotherapy), provide a mechanistic rationale for combining RP1 with PD-1 blockade, and present a targetable, plastic population for subsequent combinatorial strategies.

In summary, intratumoural oncolytic virotherapy with RP1 induces systemic reorganisation of the CD8^⁺^ T cell landscape, promotes persistence of precursor states with ongoing antigen engagement within the TME, increases clonal diversity across tumour sites, and re-establishes the immunological conditions required for successful checkpoint blockade.

These findings provide a mechanistic basis for the systemic clinical activity observed following local OV administration and support a model in which viral therapy primes or restores a precursor CD8^⁺^ T cell axis that is central to durable anti-tumour immunity in patients with ICI-resistant melanoma.

## Methods

### Mice and tumour models

Nr4a3-Tocky^15^ and Kaede transgenic mice^16^ were maintained on a C57BL/6 background. Nr4a3-Tocky mice carry a BAC transgene in which the first coding exon of *Nr4a3* was replaced with the Fluorescent Timer gene (FTfast) by knock-in/knock-out strategy and were crossed with Foxp3-IRES-GFP (C.Cg-Foxp3tm2Tch/J). Kaede mice were a kind gift from the David Withers laboratory. Female C57BL/6J, Nr4a3-Tocky, or Kaede mice (8–10 weeks) were housed under specific pathogen-free conditions with institutional ethical approval. All experiments were approved by the Animal Welfare and Ethical Review Body at the Institute of Cancer Research and conducted under UK Home Office regulations (Animals (Scientific Procedures) Act 1986). The BRAF^V600E^ 4434 melanoma line^20^ was cultured in DMEM plus 10% FBS and implanted subcutaneously on each flank (4×10^6^ cells per flank). When tumours reached ∼50 mm^3^, the right-sided tumour received intratumoural RP1 (1×10^6^ PFU in 50 μL PBS) or vehicle. Three doses were delivered on alternate days and tumour volume and body weight recorded three times weekly.

### RP1 oncolytic virus

RP1 is an HSV-1 OV engineered from strain RH018A^4^ with deletions of ICP34.5 and ICP47 and insertion of murine GM-CSF and GALV-GP-R^⁻^. RP1 (1×10^6^ PFU, 50 μL) was delivered intratumourally to the right-sided tumour using an insulin needle under isoflurane anaesthesia every other day for 3 doses (Figs 1–5f) or 2 doses (Fig. 5g–m, to capture early migration in the Kaede experiments).

### Flow cytometry and FACS

Tumours were mechanically dissociated and digested with collagenase IV/DNase I/lipase/trypsin. Wild-type tumours were digested at 37 °C for 30 min; Tocky tumours were digested at 37 °C for 10 min then 20 min at room temperature to limit timer maturation. Lymph nodes were mashed through a 70 μm filter; spleens underwent an additional red blood cell lysis step. Samples were stained with antibodies against TCRβ, CD4, CD8, FoxP3, CD44, CD62L, KLRG1, PD-1, CD25, and ICOS with zombie aqua viability dye. Data were acquired on a Cytek Aurora and analysed in FlowJo v10. For FACS sorting, tumours from Tocky mice were processed for flow cytometry and stained with antibodies against TCRβ and far red viability dye, and TCRβ+ live cells were sorted using the FACS Aria into 4 Tocky quadrants (New, Persistent, Arrested and Timer negative) prior to labelling for scRNAseq.

### Spatial profiling

Multiplex immunofluorescence (Akoya) was performed on FFPE sections using antibodies to CD4, CD8, F4/80, podoplanin, and DAPI. Images were acquired on Vectra Polaris and analysed with inForm to quantify spatial immune neighbourhoods.

### Cytokine measurements

Tumours were homogenised in lysis buffer using Precellys lysing tubes (Bertin Instruments) and lysates clarified by centrifugation. Cytokine concentrations were quantified using the LEGENDplex Mouse Cytokine Panel (BioLegend) and normalised to tumour weight.

### Tocky analysis

Tocky reporter fluorescence was assessed by spectral flow cytometry on a Cytek Aurora with spectral unmixing from single-stained controls. Timer angle transformation and Tocky locus analysis were performed as previously described^15^. Unsupervised clustering and UMAP projection used the CRAN package umap; k-means clustering on PCA output with optimal cluster number determined by Bayesian information criterion using mclust. Plots were generated with ggplot2.

### Single-cell RNA sequencing

CD8^⁺^ T cells were sorted into Tocky-defined populations and barcode-labelled (BD Rhapsody). Single-cell libraries for whole-transcriptome analysis and VDJ TCR capture were generated using BD Rhapsody reagents (BD Biosciences) and sequencing data pre-processed using Seven Bridges Genomics (BD). Quality control, normalisation, and clustering were performed in Seurat, with complementary analyses using ProjecTILs for reference annotation, ESCAPE for pathway activity inference, Slingshotfor trajectory analysis, and scVelo for RNA velocity.

### Trajectory analysis

Pseudotime trajectories were reconstructed using Slingshot on highly variable genes from normalised expression matrices. RNA velocity was estimated using scVelo with dynamical modelling and visualised by projection onto UMAP embeddings.

### Kaede photoconversion

Subcutaneous tumours (∼50–70 mm^3^) were photoconverted under isoflurane anaesthesia using a 405-nm LED light source (Dymax BlueWave Visicure) with a light-shielding guard, using nine sequential illumination cycles as previously described^17^. Tissues were harvested 48 h after photoconversion for analysis by flow cytometry.

### Cross-species gene signature analysis

Murine Tocky-defined gene signatures were derived from scRNA-seq data using Seurat FindMarkers and converted to human orthologues by one-to-one orthology mapping. Signatures were applied to bulk RNA-seq data from IGNYTE and publicly available melanoma ICI datasets. Enrichment was quantified using GSVA and singscore; comparisons between responders and non-responders used two-sided Wilcoxon rank-sum tests.

### Statistics

In vivo data are presented as mean ± s.e.m. Two-group comparisons used unpaired two-tailed Student’s t-tests; multi-group comparisons used one-way ANOVA with post hoc correction. Survival curves were analysed by log-rank (Mantel–Cox) test. P < 0.05 was considered statistically significant. Analyses were performed in GraphPad Prism.

## Data availability

scRNA-seq datasets will be deposited in the Gene Expression Omnibus (GEO) prior to publication. All other data are available from the corresponding author upon reasonable request.

## Acknowledgements

This work was supported by Cancer Research UK and the British Skin Foundation. We thank the flow cytometry facilities at the Institute of Cancer Research and Imperial College London. We acknowledge Replimune for providing RP1, grant support for experimental studies, and access to IGNYTE sequencing data, and we are grateful to the study participants. We acknowledge support from the ICR/RMH NIHR Biomedical Research Centre.

## Author contributions

E.A., A.M., K.H., and M.O. conceived the study. E.A., S.F., V.R., and J.K. performed murine experiments and flow cytometry. E.A. and J.H. performed Tocky analyses with guidance from M.O. E.A., A.R., V.R., and L.H. collected scRNA-seq data. E.A. and A.P. performed scRNA-seq analyses with guidance from M.O. E.A. wrote the manuscript with input from all authors and senior authorship from A.M., K.H., and M.O.

## Supplementary Figure Legends

**Supplementary Fig. 1.**
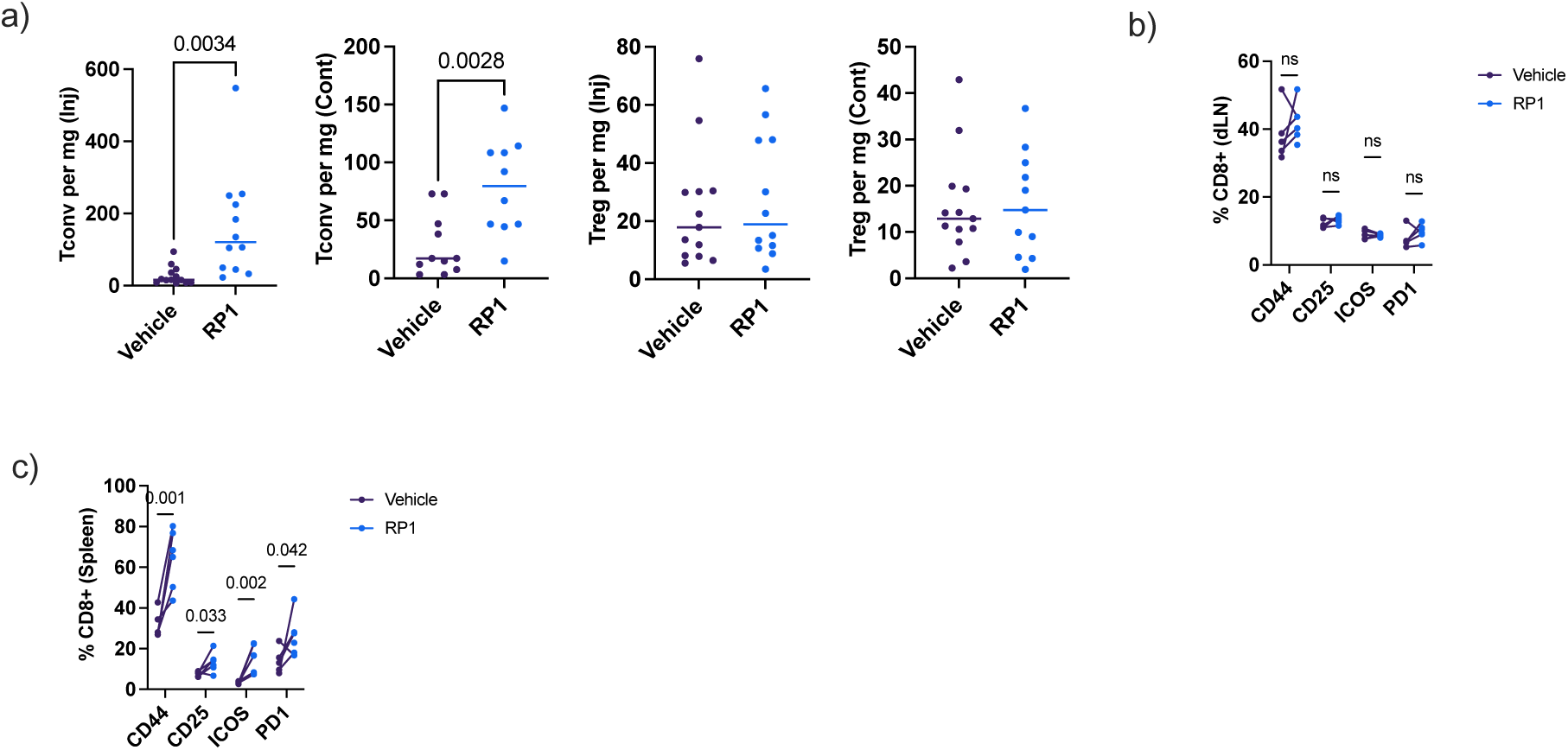
Additional flow cytometry from the RP1-treated bi-flank melanoma model. **a,** RP1 significantly increased FoxP3^⁻^ CD4^⁺^ T conventional cell infiltration at both tumour sites. **b,** No significant change in CD8^⁺^ T cell activation markers in tumour-draining lymph nodes. **c,** Significant upregulation of CD44, CD25, PD-1, and ICOS in splenic CD8^⁺^ T cells from RP1-treated mice.

**Supplementary Fig. 2.**
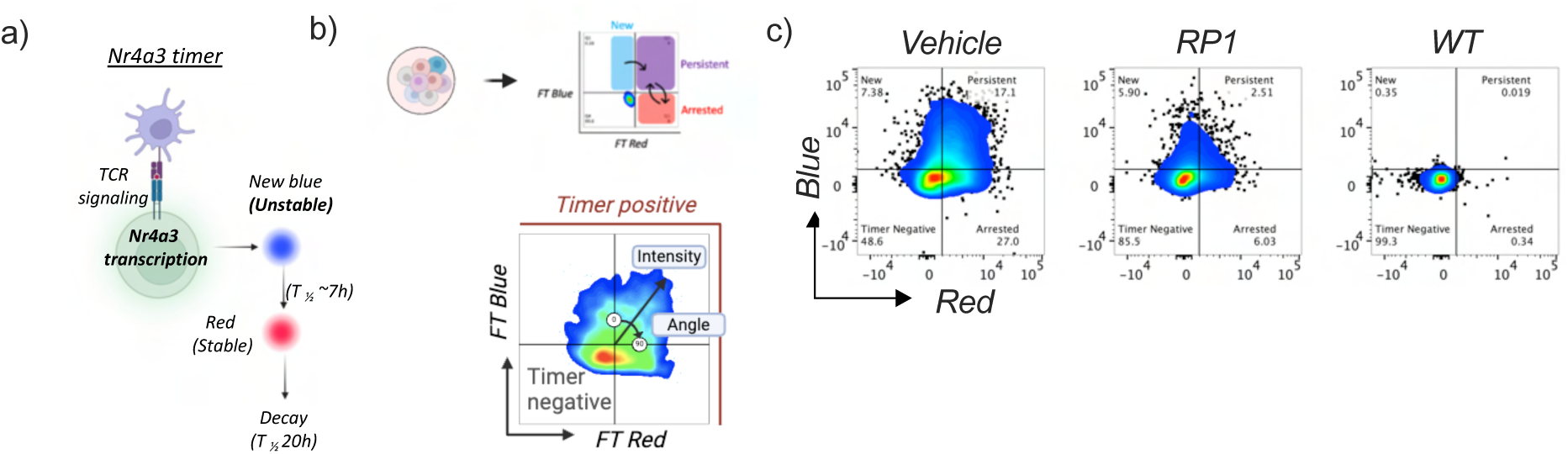
Nr4a3-Tocky reporter system and representative gating. **a,** TCR engagement drives expression of an unstable blue protein that matures irreversibly to red (t½ ∼4 h). Quadrants: blue^⁺^ = new TCR signal; blue^⁺^red^⁺^ = sustained signalling; red^⁺^ = ceased signalling; timer-negative = no TCR activation. **b,** Timer angle versus intensity gating. **c,** Representative Tocky flow plots from vehicle-treated, RP1-treated, and wild-type tumour controls.

**Supplementary Fig. 3.**
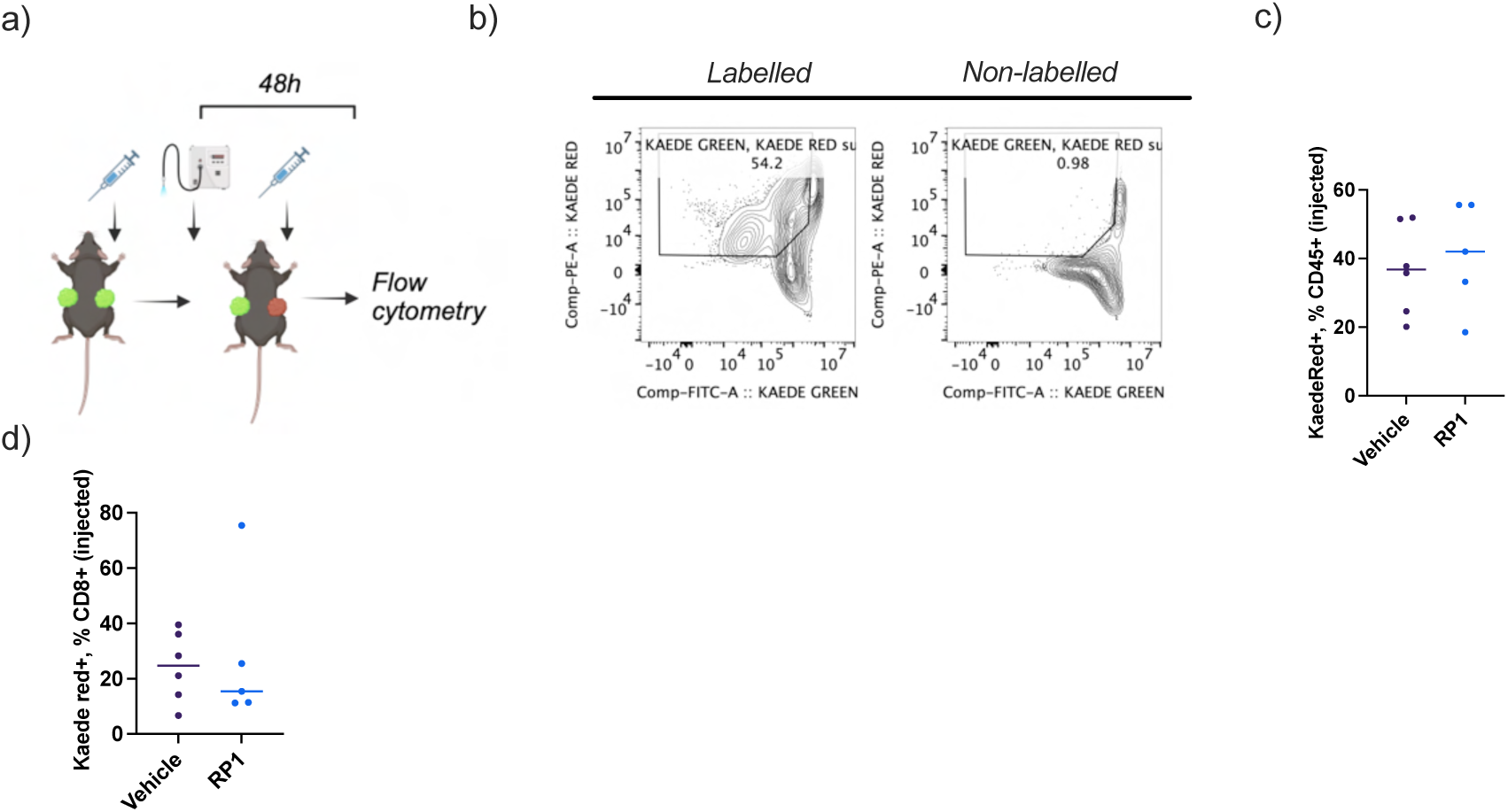
Kaede photoconversion of intratumoural T cells in vivo. **a,** Schematic of transdermal UV photolabelling. **b,** Representative Kaede red/green gating from labelled and unlabelled tumours. **c,d,** At 48 h post-photolabelling, approximately 40% of CD45^⁺^ cells (c) and 25% of CD8^⁺^ T cells (d) within the injected tumour were Kaede red.

**Supplementary Fig. 4.**
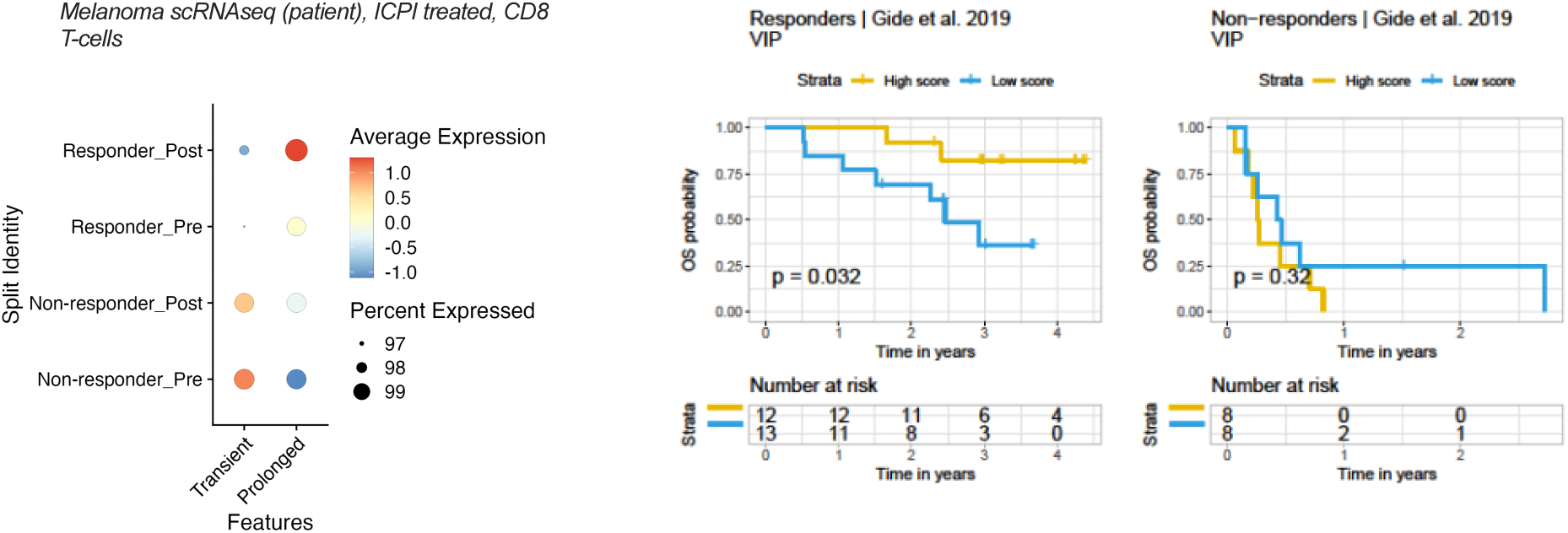
ProjecTILs reference annotation reveals distinction between viral and tumour Tpex states induced by RP1. **a,** Tocky-sorted CD8^⁺^ T clls mapped onto combined TIL and LCMV reference atlases; vehicle cells mapped to Tex populations, RP1-treated cells to naïve/Tpex (TIL reference) and viral Tpex (LCMV reference). **b,** Cluster proportions by TIL annotation. **c,** Differential expression comparing viral and TIL Tpex: increased *Tcf7* and *Slamf6* in viral Tpex, hallmarks of the VIP population.

**Supplementary Fig. 5 VIP signature predicts response to ICI in independent melanoma cohorts.**

**a,** Enrichment of VIP but not VITE programmes in CD8^⁺^ T cells from responding tumours in publicly available scRNA-seq data from patients receiving primary ICI. **b,** Kaplan–Meier analysis of the GIDE melanoma cohort: high VIP score associated with improved overall survival in ICI responders.

